# Sulforaphane modulates microRNA expression in colon cancer cells to implicate the regulation of oncogenes CDC25A, HMGA2 and MYC

**DOI:** 10.1101/183475

**Authors:** Christopher A. Dacosta, Claudia Paicu, Irina Mohorianu, Wei Wang, Ping Xu, Tamas Dalmay, Yongping Bao

**Affiliations:** Norwich Medical School, University of East Anglia, Norwich, NR4 7UQ, UK; School of Computing Sciences, University of East Anglia, Norwich, NR4 7TJ, UK; School of Biological Sciences, University of East Anglia, Norwich, NR4 7TJ, UK

**Keywords:** sulforaphane, microRNA (miRNA), cancer, colorectal, cruciferous vegetable

## Abstract

Colorectal cancer is an increasingly important cause of morbidity and mortality, whose incidence is associated with dietary and lifestyle factors, particularly inversely so with the consumption of cruciferous vegetables. These vegetables contain glucosinolates, from the breakdown of which are derived isothiocyanates, such as sulforaphane. Sulforaphane is well-characterised for wide-ranging tumour-suppressive and chemoprotective activities *in vitro*, yet deeper elucidation of its biological interactions would aid in better realising its potential in chemoprevention and/or chemotherapy. There is evidence to suggest that sulforaphane modulates microRNA expression in the colon, thus implying the potential for microRNA modulation to play a role in the anti-cancer effects of sulforaphane. Therefore, the effects of sulforaphane on microRNA expression profiles in the colonic adenocarcinoma Caco-2 and non-cancerous colonic CCD-841 cell lines were investigated by small RNA cloning and deep sequencing, followed by Northern Blot validation experiments. Sulforaphane upregulated let-7f-5p and let-7g-5p expression at 24 h in Caco-2 cells, but not in CCD-841. Such treatment also downregulated miR-29b-3p in Caco-2. Dual luciferase assays with a let-7f-5p mimic and inhibitor confirmed the binding of the miRNA to predicted binding sites in the mRNA transcript 3’-UTRs of cell division cycle 25A (CDC25A), high-mobility group AT-hook-2 (HMGA2) and MYC. Therefore, we hypothesize that let-7f-5p translationally represses CDC25A, HMGA2 and MYC, thereby playing a role in the tumour-suppressive effects of sulforaphane. The apparent selectivity of let-7f-5p induction towards tumour cells would be therapeutically desirable if applicable *in vivo*. MiR-29b-3p is predicted to target a number of tumour-suppressing genes, further investigation of which could be informative regarding the potential of sulforaphane to suppress tumour progression.

## Introduction

Colorectal cancer is an increasingly major cause of global morbidity and mortality, the latter of which increased by 46% between 1990 and 2010 (1). Its global incidence was 1.4 million in 2012 (2), making it the third most commonly diagnosed type of cancer (3). It originates from the oncogenic transformation of epithelial cells lining the colon or rectum, which are particularly susceptible to the oxidative, mutagenic and/or inflammatory effects of ingested compounds or products of the gut microbiota. Therefore, it is unsurprising that its incidence has been repeatedly linked to lifestyle and dietary factors, and observed to rise with the adoption of ‘Western’ dietary habits in a given country (4). Preventative strategies and the screening of asymptomatic individuals for early-stage colorectal cancer are vital, because noticeable symptoms tend not to arise until the cancer has progressed to an advanced and often terminal stage (5).

Colorectal cancer risk is particularly inversely correlated with cruciferous vegetable consumption (6), which is likely to be attributable to the derivation of isothiocyanates (ITCs) from the glucosinolates: a group of compounds particular to this group of vegetables that includes broccoli, Brussel sprouts, cabbage and bok choy. Plant cell rupture (e.g. by cooking and/or chewing) enables myrosinase enzymes to hydrolyse the glucosinolates to form several types of product, including ITCs. Broccoli is high in a particular glucosinolate, glucoraphanin, from which is derived a specific ITC known as sulforaphane (SFN). SFN, amongst other ITCs, is a well-established inducer of the nuclear factor (erythroid-derived 2)-like 2 (Nrf2) transcription factor, which upregulates the transcription of various antioxidant genes whose promoters contain the *antioxidant response element*. These help to protect against potentially oncogenic, oxidation-induced, mutagenesis, such as by inhibiting 2-amino-1-methyl-6-phenylimidazo[4,5-b]pyridine (PhIP)-induced DNA damage (7). It is also widely reported to directly suppress oncogenic characteristics of tumour cell lines, such as cell migration, invasion and proliferation, and to promote cell cycle arrest and apoptosis in such (8).

Moreover, SFN is a hormetic agent, in that the medium-term consequences of low-to-moderate exposure - i.e. that reasonably achievable in humans via vegetable consumption - tend to be cytoprotective, whereas much higher-dose exposure can be cytotoxic via the induction of oxidative stress and DNA damage (9). This is because SFN acts acutely as a prooxidant via several mechanisms, including the depletion of cellular glutathione (10) and the disruption of the mitochondrial respiratory chain (11). Low-to-moderate SFN exposure can trigger an Nrf2-mediated anti-oxidative cellular response whose reductive effects outweigh those of the initial pro-oxidant effect in the medium term. On the other hand, the pro-oxidant effects of high-dose exposure may be such as to induce significant oxidative damage to macromolecules and thus result in cytotoxicity rather than cytoprotection. The latter form of activity is obviously undesirable if occurring in healthy cells, but inevitably contributes to the tumour-suppressive effects of SFN observed in tumour cell lines, as described above. Conversely, while the cytoprotective capacity of low-to-moderate dose SFN is desirable in normal cells and helps to protect them against oncogenic transformation, such cytoprotection of tumour cells is undesirable as it may increase their resistance to cytotoxicity-dependent forms of anti-cancer therapy. This has led to speculation that low-to-moderate doses of SFN, whether dietary or supplemental, might even be detrimental to patients undergoing treatment for advanced cancers. However, the ‘cytotoxic threshold’ of SFN with respect to dose inevitably varies according to cell type, due to variations in basal levels of reactive oxygen species and cell signalling. Notably, a number of studies have indicated tumour cell lines to be more sensitive than their non-cancerous counterparts to SFN-induced cytotoxicity (i.e. to have a lower ‘cytotoxic threshold’) (12), which opens up the possibility of using SFN at intermediate doses that are simultaneously cytotoxic towards tumour cells yet cytoprotective towards surrounding non-cancerous cells.

Clearly, the bioactivity of SFN is highly complex and far from fully understood. Further understanding of SFN’s biological interactions is crucial in order to further investigate its potential chemopreventative and/or chemotherapeutic uses. There is evidence to suggest that SFN is able to modulate the expression of miRNAs in several colonic cell lines, from a study by Slaby et al. (13). Given that miRNAs are well-established as major players in human development and disease progression via their post-transcriptional regulation of gene expression, it is highly probable that SFN’s anti-cancer effects are partly attributable to the modulation of miRNA expression. Therefore, a study to investigate the modulation of miRNA expression by SFN in a colonic adenocarcinoma cell line, Caco-2, and a noncancerous colonic cell line, CCD-841, was carried out by first performing a wide-scale assay of SFN-mediated miRNA modulation by small RNA (sRNA) library cloning and deep sequencing, and then verifying the SFN-mediated modulation of specific miRNAs by individual assays. Subsequently, the ability of modulated miRNAs to bind to several computationally-predicted oncogenic mRNA targets was verified *in vitro*. The insight into the potential bioactivity of SFN provided by the findings of the study with respect to colorectal carcinogenesis could lead to new avenues of research regarding the potential applications of diet-derived and/or supplemental SFN in chemoprevention, and/or standalone or adjuvant chemotherapy.

## Materials and Methods

### Materials

SFN (1-isothiocyanato-4-[methylsulfinyl]-butane, 99% purity (Cat No. S699115) was purchased from Toronto Research Chemicals (Canada). Dimethyl sulfoxide (DMSO), Dulbecco’s Modified Eagle’s Medium containing D-glucose (4.5g/L) and non-essential amino acids (DMEM) (Cat. no. D5671-500ML), heat-inactivated FBS (Cat. no. 12106C-500ML), L-glutamine (Cat. no. 59202C-100ML), Penicillin-Streptomycin (Cat. no. P4083-100ML), DNA oligonucleotides, TRI Reagent (Cat. no. T9424), the mirPremier™ MicroRNA Isolation Kit (Cat. no. SNC50), and the PerfectHyb™ Plus Hybridization Buffer (Cat no. H7033) were all purchased from Sigma-Aldrich (Dorset, UK). The let-7f-5p mimic (Cat. no. MSY0000067; target sequence: 5’-UGAGGUAGUAGAUUGUAUAGUU-3’), let-7f-5p inhibitor (Cat. no. MIN0000067; target sequence: 5’-UGAGGUAGUAGAUUGUAUAGUU-3’), AllStars Negative Control siRNA (Cat. no. SI03650318), and Attractene Transfection Reagent (Cat. no. 301005) were all purchased from Qiagen (West Sussex, UK). RNAse Inhibitor (Cat. no. AM2682), Gel Loading Buffer II (Cat no. AM8546G), T4 Polynucleotide Kinase Forward Reaction Buffer (Cat. no. 10343010) and T4 Polynucleotide Kinase (Cat. no. EK0032) were all purchased from Fisher Scientific (Loughborough, UK). The Zymo RNA Clean & Concentrator™ -25 kit (Cat. no. R1017) was purchased from Cambridge Bioscience (Cambridge, UK). The Dual-Glo^®^ Luciferase Assay System (Cat. no. E2920) was purchased from Promega UK (Southampton, UK). The human colorectal adenocarcinoma Caco-2 (HTB-37) and human normal colonic CCD-841 (CRL-1790) cell lines were obtained from ATCC.

### Cell Culture and Treatments

Cells were maintained in DMEM supplemented with FBS (10%) (v/v), L-glutamine (200nM), and Penicillin-Streptomycin. Cells were cultured at 37°C in air under 5% CO2. Caco-2 cells were split before exceeding 80% confluence, to avoid their differentiation. Caco-2 cells were tested in the laboratory and found to be mycoplasma free. CCD-841 cells were not tested for mycoplasma in the laboratory.

SFN was dissolved in DMSO, and final DMSO concentrations were kept at 0.05%. For sRNA library construction and Northern Blotting experiments, Caco-2 cells were cultured to 70-80% confluence in 10cm dishes, then treated in duplicate with SFN (10µM) in culture medium for 8 or 24 h, or DMSO (control). For luciferase assays, HCT-116 cells were subject to transfection with pmiRGLO vectors and miRNA mimic and/or inhibitor at the same time as the seeding of 3.0 × 10^4^ cells/well in 96-well plates. Co-transfection of let-7f-5p mimic and/or let-7f-5p inhibitor and/or AllStars Negative Control siRNA was performed using Attractene Transfection Reagent, according to the manufacturer’s instructions. Briefly, transfection mixtures were prepared by combining Attractene, pmiRGLO vector, and miRNA mimic (100nM) and/or inhibitor (100nM) and/or AllStars Negative Control siRNA (0200nM), in non-supplemented DMEM, adjusting the amounts of AllStars Negative Control siRNA in order to maintain a 200nM RNA concentration. These mixtures were incubated at room temperature for 15 min, and then added directly into suspended cells immediately following the seeding of cells into 96-well plates. Cells were subsequently incubated for 48 h.

### RNA Extraction

For sRNA library construction, total RNA was extracted using TRI Reagent, according to the manufacturer’s instructions (with minor modifications to the protocol). Briefly, cells were lysed in TRI Reagent by directly adding reagent onto attached cells for ≥ 1 min following the removal of culture medium. Lysates were incubated for 5 min at room temperature, then 200µL chloroform was mixed per 1mL lysate. Mixtures were incubated at room temperature for 10 min after vigorous shaking, then centrifuged at 16 000 *g* for 15 min at 4°C. The resulting aqueous, uppermost phases were retained, from which RNA was then recovered by isopropanol precipitation (≥5 h at -20°C) and then centrifugation at ≥ 16 000 *g* for 30 min. Pellets were recovered, washed in 75% ethanol, dried, and then resuspended in nuclease-free H_2_O with RNAse Inhibitor.

For Northern Blot experiments, either total RNA was extracted in the same manner as that described above, or small RNA was extracted using the mirPremier™ MicroRNA Isolation Kit according to the manufacturer’s instructions.

### Small RNA Library Construction

Total RNA samples were integrity-checked by agarose gel electrophoresis to visualise 28S and 18S bands, and then purified using the Zymo RNA Clean & Concentrator™ -25 kit according to the manufacturer’s instructions. Subsequently, sRNA libraries were constructed from the purified total RNA samples according to the protocol published in 2015 by Xu et al. (14) with the HD adapters previously developed by Sorefan et al. (15), using 2µg total RNA per library.

### Small RNA Library Sequencing and Bioinformatics Analysis

Nine indexed sRNA libraries were multiplexed on a sequencing lane and deep-sequenced on the Illumina HiSeq 2500 platform by the Earlham Institute, formerly known as The Genome Analysis Centre (Norwich, UK), with a sequencing depth of 100 million reads per lane.

Analysis of the deep sequencing data was performed using custom-made Perl and R scripts. Firstly, the adapters were trimmed based on a perfect match to the first 8 nt of the adapter sequence (TGGAATTC). Next, the HD signatures (assigned *NNNN* fragments localized at the 5’ and 3’ ends of each read) were trimmed. The reads were converted into non-redundant format. The unique sequences and their abundances i.e. the number of occurrences in a sample, were logged. The reads were then mapped to the reference human genome *(H. sapiens* v. 38 (16)), the available annotations, and the *H. sapiens* miRNAs downloaded from miRBase v. 21 (17), using PatMaN (18), allowing 0 gaps and ≤ 2 mismatches (see Supplementary Table S1 for a summary of raw and accepted read numbers per library). Next, the read abundances were normalized by the reads per million (RPM) (19), sRNA-adapted quantile normalization (20) and subsampling (21) methods. The most appropriate normalization method for this dataset was determined to be RPM, conducted on a scaling total of 13 000 000: the median total value. The differential expression analysis was done on the median expression levels of the biological replicates as pairwise comparisons, using an offset fold-change approach (offset=20) (22). All miRNAs with a fold-change ≥ 1.9 between either control and 8 h, control and 24 h, or 8 h and 24 h, were predicted as differentially-expressed candidates.

The data corresponding to this study are publicly available in the Gene Expression Omnibus (GEO) (23) under accession number GSE89363 (samples GSM2367444 to GSM2367459). For library size class distributions, see Supplementary Figure S1. For complexity distributions, see Supplementary Figure S2. For the read annotation analysis and determination of differential expression, see Supplementary Tables S2, S3 and S4.

### Northern Blotting

Equal amounts of total RNA (10µg) or small RNA (5µg) were combined with Gel Loading Buffer II, denatured at 65°C for 5 min, then size-fractionated by urea-PAGE in pre-run 16% gels at 100V for 2 h, in 0.5X tris/borate/EDTA buffer (TBE). Size-fractionated RNA was transferred to Hybond^®^ NX membranes by semi-dry electroblotting in 0.5X TBE at 3mA/cm^2^ for 1 h. RNA was then chemically cross-linked to membranes using a solution prepared by diluting 0.373g 1-ethyl-3-(dimethylaminopropyl) carobiimide in 10mL dH_2_O, then adding 122.5µL 12.5M 1-methylimidazole and 10µL 1M HCl. Membranes were placed face-up onto filter paper soaked with the solution, sealed in Saran wrap, then incubated at 60°C for 1 - 2 h. Following cross-linking, membranes were washed in dH_2_O for 10 min.

PerfectHyb™ Plus hybridization buffer was pre-incubated at 37°C to dissolve precipitates, then membranes were incubated in hybridization buffer at 37°C under rotation for 1 h. Radioactive miRNA probes were prepared by mixing T4 Polynucleotide Kinase Forward Reaction Buffer, DNA oligonucleotide antisense to the miRNA of interest (or part of human U6) (10µM), T4 Polynucleotide Kinase, dH_2_O, and γ-ATP (P-32), then incubating at 37°C for 1 h. Prepared radioactive probes were added to hybridization buffer, in which membranes were subsequently incubated at 37°C under rotation overnight. Then, membranes were washed four times in 0.2X SSC/0.1% SDS solution and sealed in Saran wrap. Phosphorimaging plates were exposed to the radioactively-labelled membranes for appropriate periods of time (typically 6-24 h), and then signals were evaluated using a phosphorimaging scanner. RNA band intensities were densitometrically analyzed using the ImageJ software. Prior to re-probing for different targets, membranes were stripped by incubation in near-boiling 0.1% SDS.

### Dot Blotting

Dot blots were prepared by adding DNA oligonucleotides of sequence sense to those of miRNAs of interest directly onto Hybond® NX membranes, then chemically cross-linking them to the membranes in the same manner that blotted RNA was cross-linked to membranes for Northern Blotting. Radioactive probe preparation, hybridization, and phosphorimaging were subsequently performed in the same manner as for Northern Blotting.

### Luciferase Reporter Assays

Dual luciferase reporter assays *(Firefly/Renilla)* were performed using the Dual-Glo^®^ Luciferase Assay System, according to the manufacturer’s instructions, and luminescence was measured using the FLUOstar Omega microplate reader (BMG Labtech), using non-transfected cells to determine background luminescence. Firefly luminescence values were normalized to *Renilla* luminescence values to control for any well-to-well variation in cell viability and/or transfection efficiency.

## Results

### Sulforaphane-Induced MicroRNA Differential Expression

The sRNA-seq data analysis indicated that the treatment of Caco-2 cells with SFN (10µM) for 8 or 24 h modulated the expression of 42 miRNAs by ≥ 1.9-fold, based on pairwise comparisons between time points of the medians of replicate normalized expression levels. Overall, let-7f-5p and let-7g-5p were the most abundant mature miRNAs, and were upregulated 1.4- and 2.1-fold by 8 and 24 h SFN treatment, respectively, as illustrated by the data presented in Figure 1. Reads for miR-193b-3p were of moderate abundance, and indicated 1.5- and 2.0-fold downregulation by 8 and 24 h SFN treatment, respectively, as illustrated by the data presented in Figure 1.

**Figure 1:**
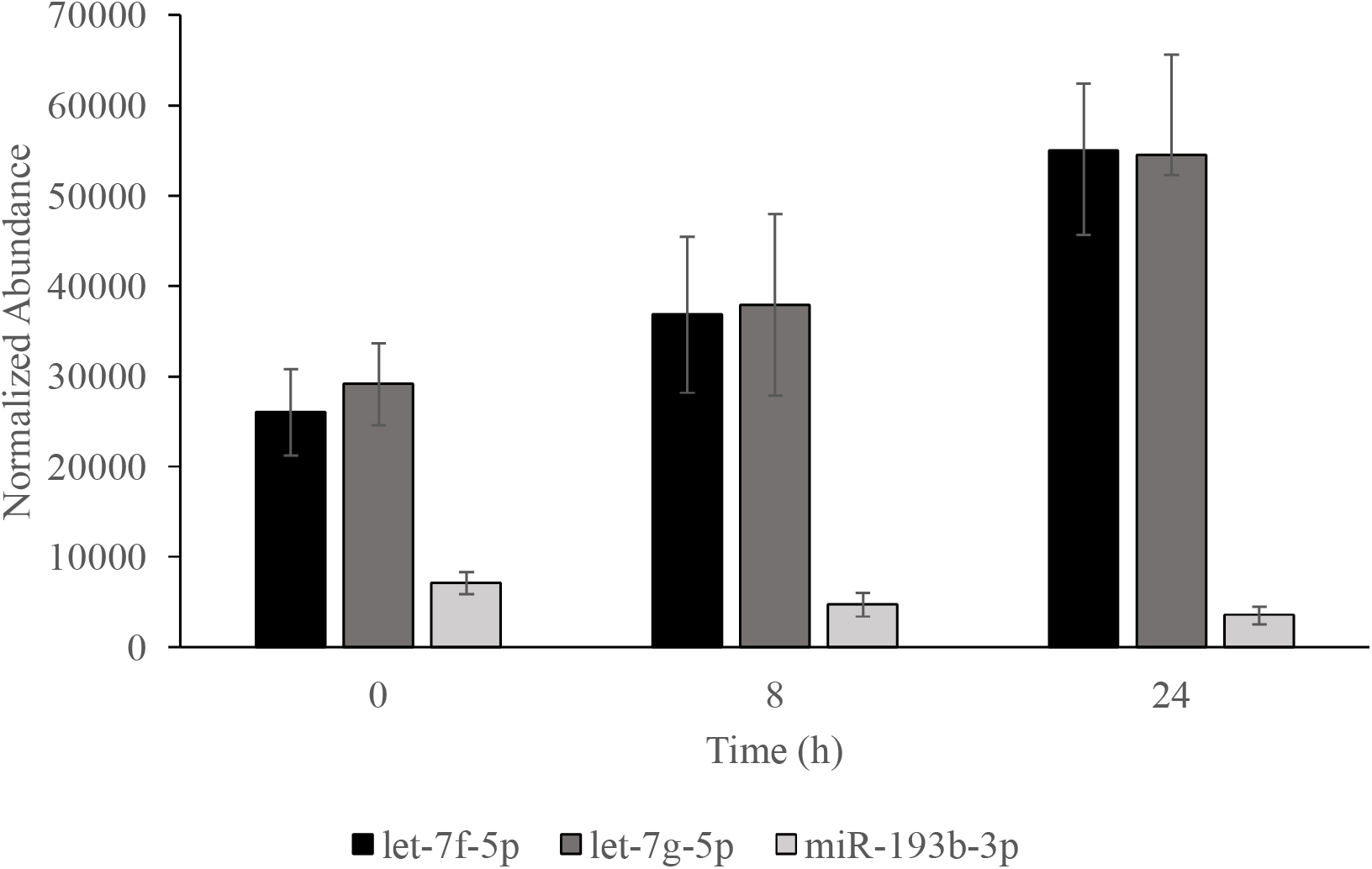
Normalized library abundances of let-7f-5p, let-7g-5p and miR-193b-3p in Caco-2 following 8 or 24 h SFN (10µM) treatment, or DMSO controls (0 h). Data are represented as the median abundance at each time point (of duplicates at 0 and 8 h, and of triplicates at 24 h), with the maximum and minimum values of replicate sets represented by the upper and lower bounds of error bars.

Northern Blot experiments confirmed that SFN (10µM) upregulated let-7f-5p and let-7g-5p expression in Caco-2 cells; the former by 1.2- and 1.4-fold at 8 and 24 h respectively, and the latter by 1.1- and 1.6-fold at 8 and 24 h respectively, as illustrated by the data presented in Figure 2 [A]. However, they showed that similar treatment of CCD-841 cells affected the expression of neither let-7f-5p nor let-7g-5p, as illustrated by the data presented in Figure 2 [B]. Meanwhile, Northern Blots for miR-193b-3p confirmed a downregulation in Caco-2 by 8 but not 24 h SFN treatment, as illustrated by the data presented in Figure 3 [A]. The Northern Blot experiment for miR-29b-3p indicated that it was downregulated in Caco-2 cells 1.5- and 1.7-fold by 8 and 24 h SFN treatment, respectively, as illustrated by the data presented in Figure 3 [B].

**Figure 2:**
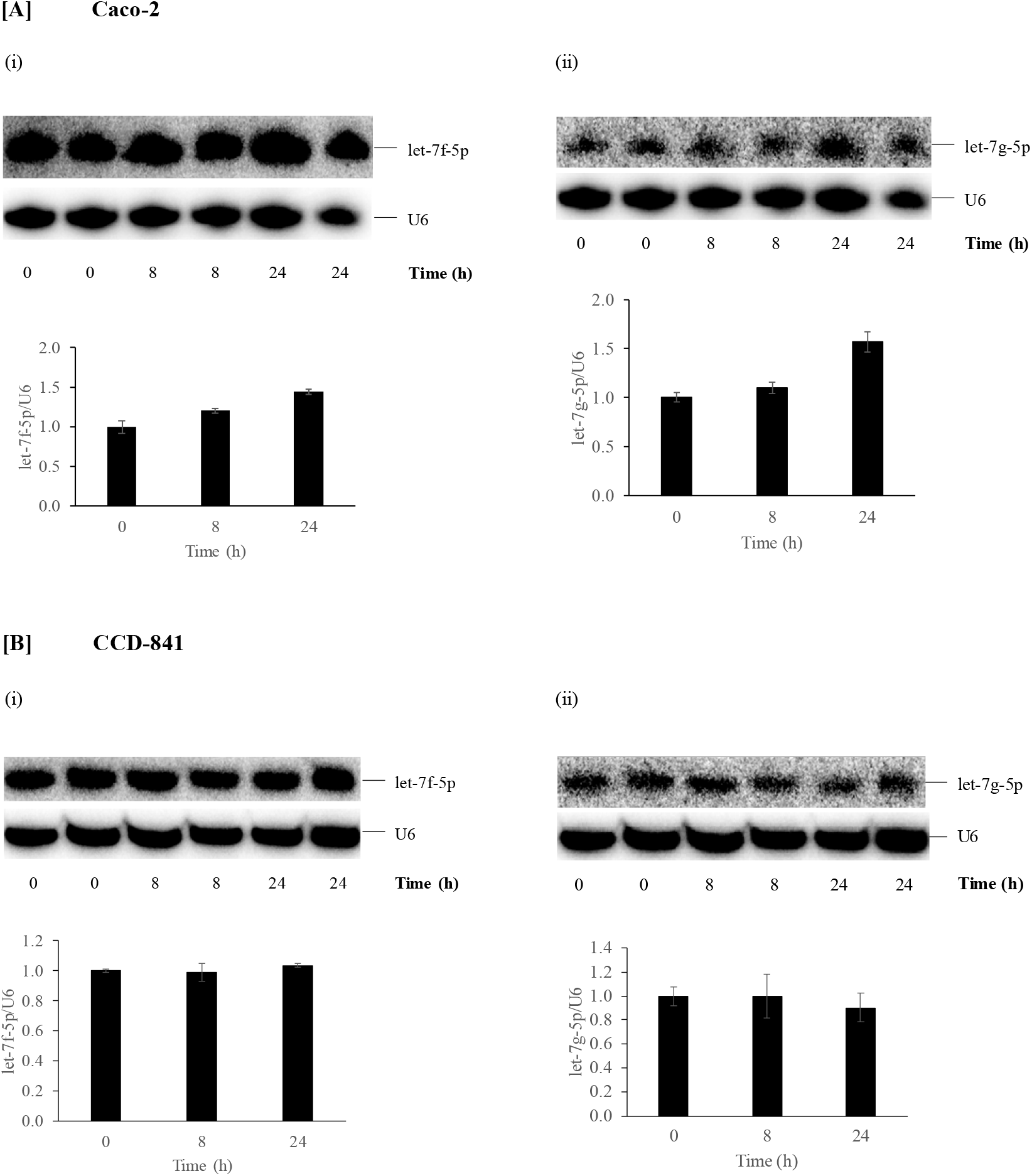
Northern Blot data for let-7f-5p and let-7g-5p in [A] Caco-2 and [B] CCD-841 cells, treated for 8 or 24 h with SFN (10μM) or DMSO controls. U6 was used as an internal control for normalization. Induction is expressed as [(miRNA band intensity)/(U6 band intensity)] for each sample relative to the control mean, and data are represented as means of duplicates ± (upper value — lower value)/2.

**Figure 3:**
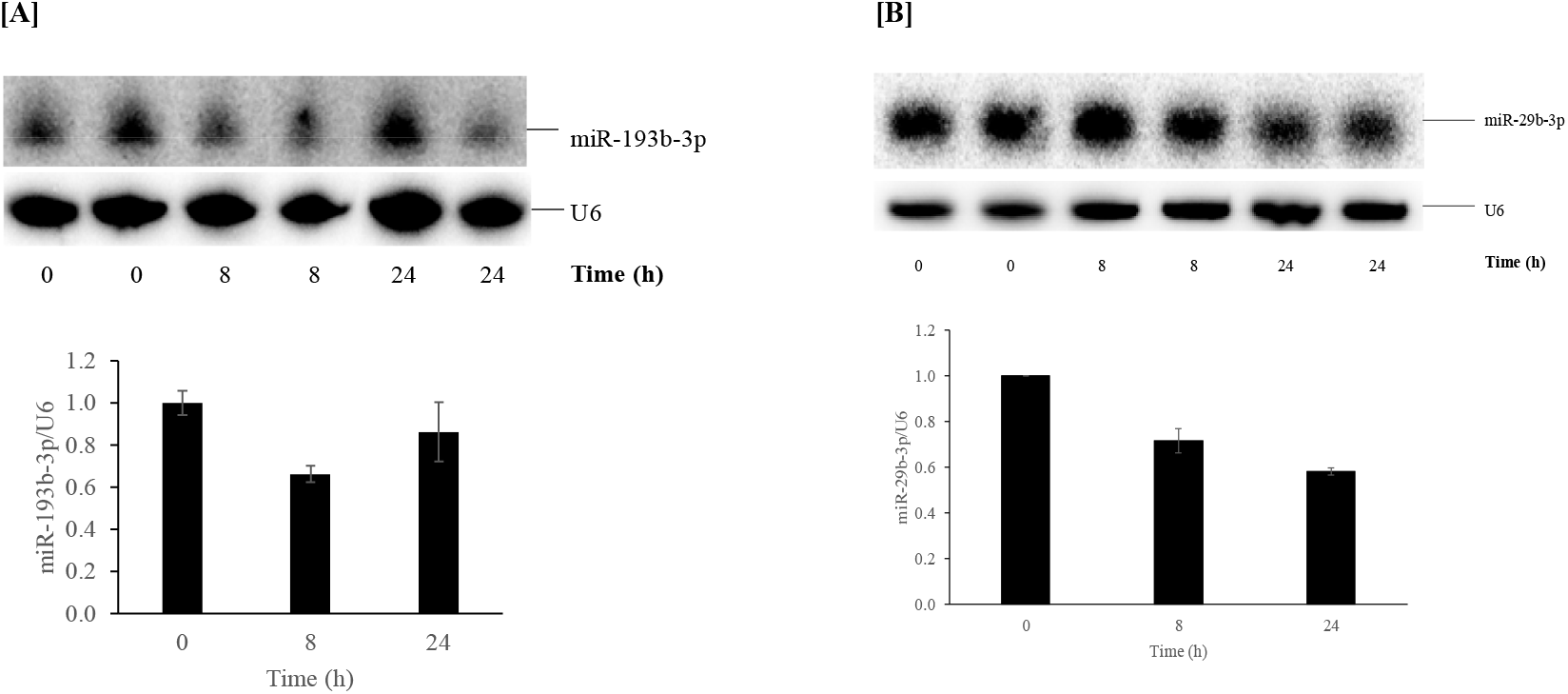
Northern Blot data for [A] miR-193b-3p and [B] miR-29b-3p in Caco-2 cells treated for 8 or 24 h with SFN (10M) or DMSO controls. U6 was used as an internal control for normalization. Induction is expressed as [(miRNA band intensity)/(U6 band intensity)] for each sample relative to the control mean, and data are represented as means of duplicates ± (upper value — lower value)/2.

### Specificity of Let-7f-5p and Let-7g-5p Northern Blot Probe Hybridization

Many miRNAs of the let-7 family are very similar in sequence, varying only at loci outside of the seed region. For example, let-7f-5p and let-7g-5p vary at positions 12 and 18, where let-7f-5p has adenine and uracil, while let-7g-5p has uracil and cytosine. This led to initial speculation that the let-7f-5p and/or let-7g-5p probe(s) used for Northern Blotting might have bound off-target to other let-7 miRNAs. In order to test the potential for such cross-hybridization, dot-blots of DNA oligonucleotides sense to the sequences of let-7a-5p, let-7b-5p, let-7c-5p, let-7d-5p, let-7e-5p, let-7f-5p, let-7g-5p and let-7i-5p were prepared, and then probed using the let-7f-5p and let-7g-5p Northern Blot probes.

Densitometric dot analysis indicated that the let-7f-5p probe was largely specific to let-7f-5p, albeit binding to let-7a-5p with affinity 6.3-fold lower than that with which it bound its target, as illustrated by the data presented in Figure 4, which also demonstrate that the let-7g-5p Northern Blot probe was largely specific for its target, albeit binding to let-7i-5p with 5.3-fold lower affinity.

**Figure 4:**
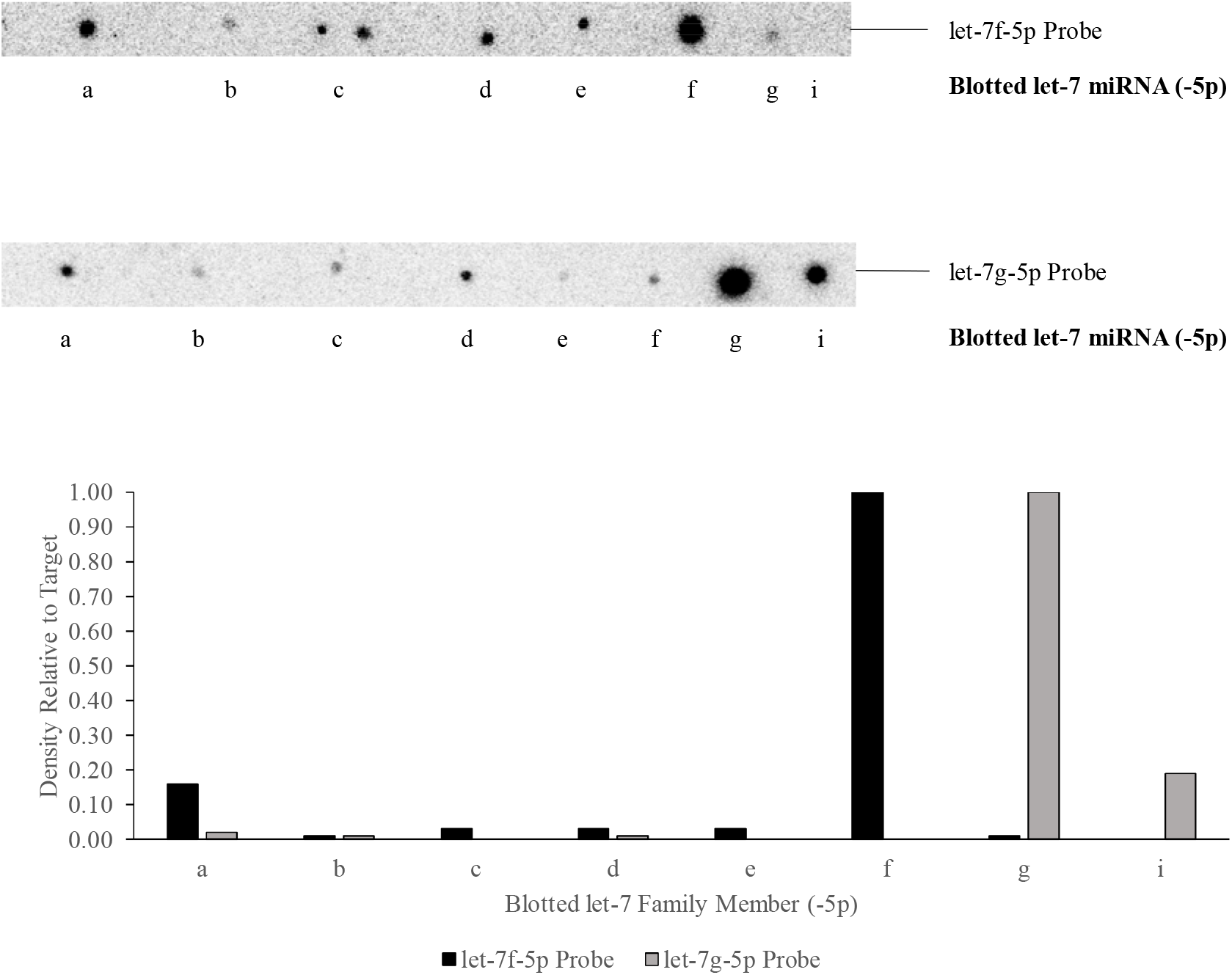
Dot Blot data following the probing of a membrane that contained positive-sense oligonucleotides representing let-7a-5p, let-7b-5p, let-7c-5p, let-7d-5p, let-7e-5p, let-7f-5p, let-7g-5p and let-7i-5p, with probes complementary to either let-7f-5p, or to let-7g-5p.

However, reads for neither let-7a-5p nor let-7i-5p were actually present in the Caco-2 sRNA library deep sequencing data.

### Targeting of mRNA 3-Untranslated Regions by Let-7f-5p

Targets of let-7f-5p predicted by the miRanda algorithm with mirSVR scores ≤ -1.2 were obtained from the online *microRNA.org - Targets and Expression* database (24), and included the mRNA transcripts of CDC25A, HMGA2, MYC and KRAS.

### Let-7f-5p Antisense-Repeats

In the luciferase assay experiments, co-transfection of the let-7f-5p mimic with the let-7f-5p antisense repeat-containing vector repressed relative Firefly luciferase expression 3.2-fold, while co-transfection of the inhibitor with the same vector upregulated relative Firefly luciferase expression 3.0-fold, as illustrated the data presented in Figure 5.

**Figure 5:**
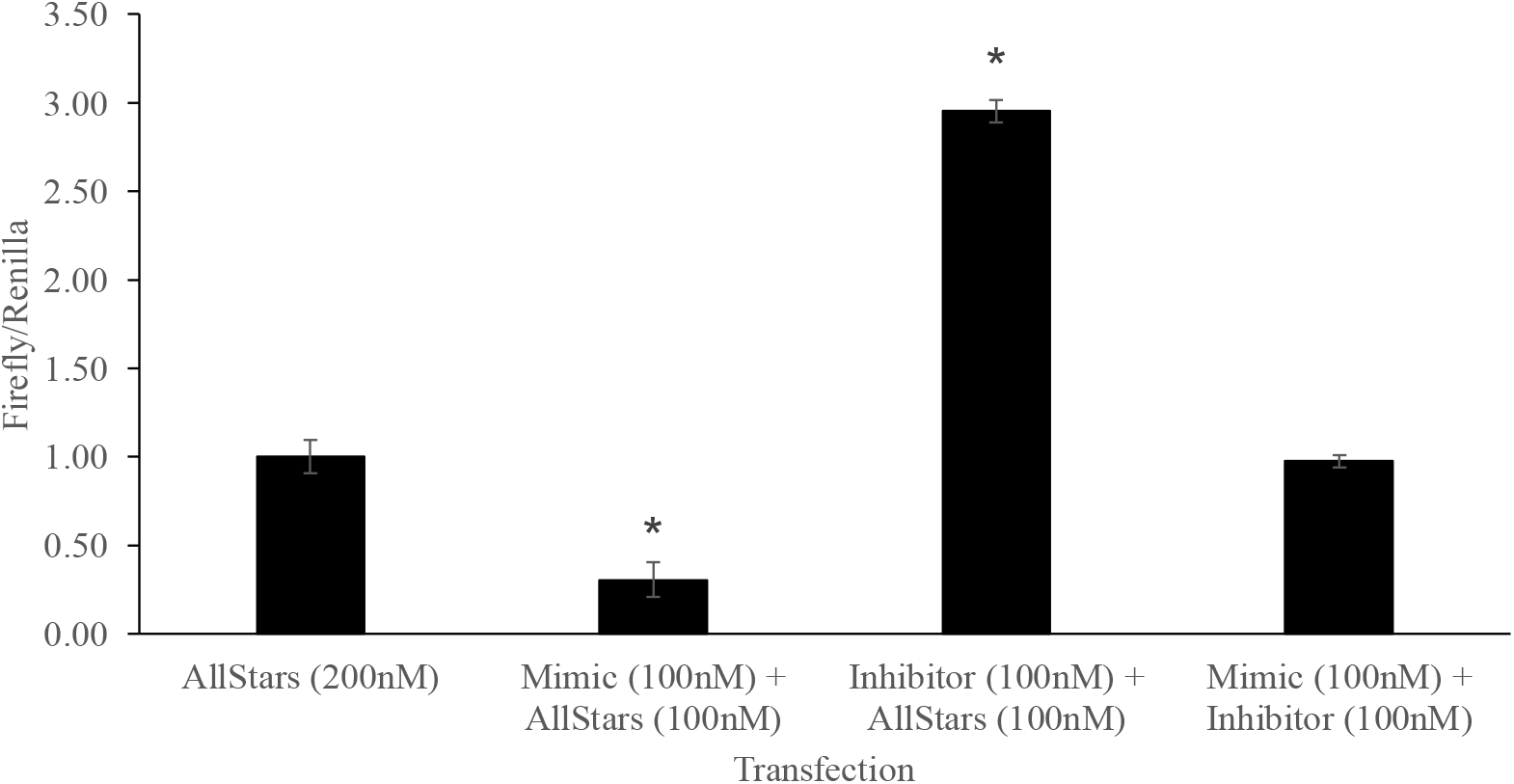
Data from luciferase assays performed 48 h after co-transfecting HCT-116 cells with pmiRGLO vector and let-7f-5p mimic and/or let-7f-5p inhibitor and/or AllStars Negative Control siRNA. The pmiRGLO vector contained an insert consisting of four repeats of sequence antisense to that of let-7f-5p. Blank-corrected Firefly luminescence values were divided by blank-corrected Renilla luminescence values, and then ratios were normalized to controls (AllStars Negative Control siRNA at 200nM). Normalized data are represented as triplicate means ± SEM (*p<0.05 according to two-tailed Student’s T-test).

Co-transfection of both the mimic and inhibitor with the vector resulted in no net modulation of relative Firefly luciferase expression.

### CDC25A, HMGA2, MYC and KRAS mRNA 3’-Untranslated Regions

Co-transfection of the let-7f-5p mimic with the vector containing part of the CDC25A 3’-UTR with a computationally-predicted let-7f-5p binding site did not affect relative Firefly luciferase expression. However, co-transfection of the let-7f-5p inhibitor with the same vector upregulated relative Firefly luciferase expression 1.4-fold, as the data presented in Figure 6 [A] demonstrate. Co-transfection of either the mimic or inhibitor with the vector containing a modified form of the CDC25A 3’-UTR portion, in which the predicted binding site was mutated, had no effect.

**Figure 6:**
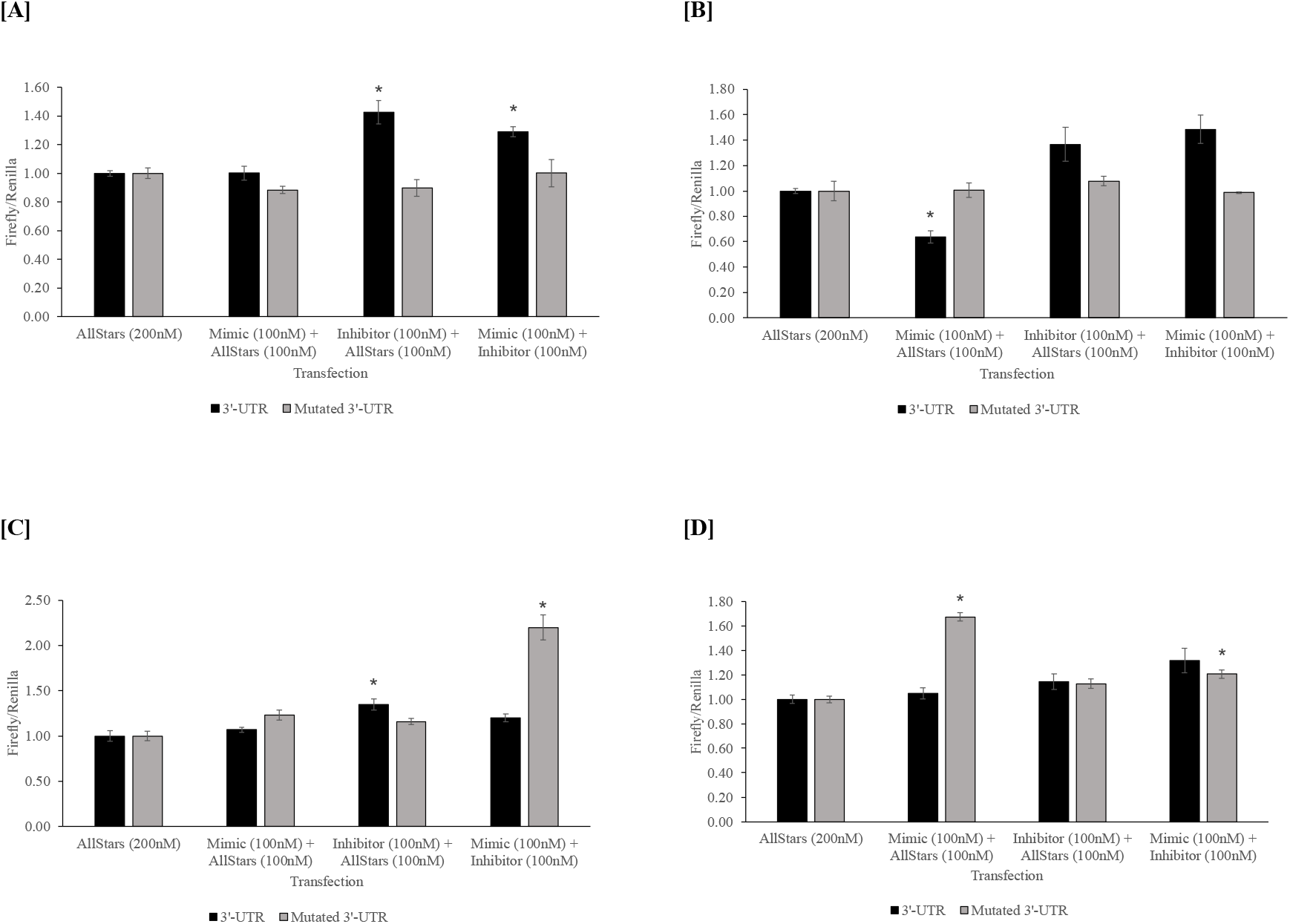
Data from luciferase assays performed 48 h after co-transfecting HCT-116 cells with pmiRGLO vector and let-7f-5p mimic and/or let-7f-5p inhibitor and/or AllStars Negative Control siRNA. The pmiRGLO vector contained an insertion of either part of the [A] CDC25A, [B] HMGA2, [C] MYC or [D] KRAS mRNA 3’- UTR containing (a) predicted let-7f-5p binding site(s), or a modified version of such in which said site(s) was/were mutated. Blank-corrected Firefly luminescence values were divided by blank-corrected Renilla luminescence values, and then ratios were normalized to controls (AllStars Negative Control siRNA at 200nM). Normalized data are represented as triplicate means ± SEM (*p<0.05 according to two-tailed Student’s T-test).

The co-transfection of the mimic with the vector containing part of the HMGA2 3’-UTR, with two predicted binding sites, repressed relative Firefly luciferase expression 1.6-fold, as the data presented in Figure 6 [B] illustrate. However, co-transfection of the inhibitor with the vector had no significant effect, but the co-transfection of inhibitor together with the mimic and vector clearly abrogated the mimic’s repressive effects. Neither the mimic nor inhibitor affected relative Firefly luciferase expression when co-transfected with the mutated-binding-site form of the vector.

Co-transfection of the mimic with the vector containing part of the MYC 3’-UTR had no effect on relative Firefly luciferase expression, but co-transfection of the inhibitor increased such expression 1.4-fold, as illustrated by the data presented in Figure 6 [C]. Neither the mimic nor inhibitor had any effect on relative Firefly luciferase expression when solely co-transfected with the mutant form of the vector, but when the mimic, inhibitor and vector were all co-transfected at once, expression from the mutant vector unexpectedly increased 2.2-fold.

The co-transfection of neither the mimic nor inhibitor had any significant effect on relative Firefly luciferase expression from the vector containing part of the KRAS 3’-UTR with one predicted binding site, as illustrated by the data presented in Figure 6 [D]. However, cotransfection of the mimic with the mutant-UTR vector unexpectedly increased relative Firefly expression 1.6-fold, while transfection of the inhibitor together with the mimic and vector partially abrogated the mimic’s effects.

## Discussion

The miRNA expression profiling data show that SFN (10µM) is able to upregulate the expression of both let-7f-5p and let-7g-5p in the colonic adenocarcinoma Caco-2 cells, but not in the non-cancerous colonic epithelial CCD-841 cells. The mechanism behind this differential effect between the cell lines is unclear. One possibility is that SFN treatment suppresses NF-ĸB in Caco-2 cells, thereby inhibiting lin28 and thus upregulating let-7 expression (25). The luciferase assay data demonstrate that let-7f-5p is indeed able to bind directly to a locus in the CDC25A 3’-UTR that was computationally predicted as a let-7f-5p binding site, and that said locus would play a role in the potential let-7f-5p-mediated translational repression of CDC25A. They also confirm that the same miRNA is able to directly bind one or more loci in the HMGA2 3’-UTR that were predicted as binding sites, and that said loci could play a role in the potential let-7f-5p-mediated translational repression of HMGA2. Finally, they suggest that the miRNA is able to directly bind to a computationally-predicted binding site in the 3’-UTR of MYC, and that this site would play a role in potential let-7f-5p-induced repression of MYC translation.

Overall, the data confirm that let-7f-5p and let-7g-5p were upregulated in Caco-2, but not in CCD-841, by SFN (10µM) treatment, and that let-7f-5p can directly interact with loci in the oncogenic CDC25A, HMGA2 and MYC 3’-UTRs. The let-7 family of miRNAs are widely reported to have anti-proliferative, apoptotic and/or cell cycle arrest-inducing effects across a number of cell types. Thus, the data presented here indicate that the translational repression of CDC25A, HMGA2 and MYC is likely to contribute to the colorectal cancer-suppressive effects of let-7 miRNAs, particularly let-7f-5p. They also demonstrate an apparent selectivity of SFN-mediated let-7f-5p and let-7g-5p induction towards the cancerous Caco-2 cell line, over the non-cancerous CCD-841 line. Such selectivity of let-7f-5p and/or let-7g-5p induction may contribute to SFN’s potential to favour cytotoxicity towards cancerous cells over that towards healthy cells as reported in several contexts, and/or may serve to counterbalance the Nrf2-mediated cytoprotective response to moderate-dose SFN that inevitably occurs in tumour cells, whilst facilitating the cytoprotection of healthy cells.

Of additional interest is the observed SFN-mediated downregulation of miR-29b-3p in Caco-2 cells. The role of miR-29b-3p appears to be ambiguous, since both oncogenic (26) and tumour-suppressive (27) functions are reported. MiR-29b-3p is computationally predicted by miRanda to target the following tumour suppressor genes (TSGs): *ADAMTS9, ARRDC3, CASP7, FEM1B, HBP1, HPGD, ING3, NAV3, PTEN, RhoBTB1, SEL1L, SLC5A8, SUV420H2, TET1, TET2* and *UVRAG* (24). Direct interactions between miR-29b-3p and the 3’-UTRs of colorectal cancer-suppressing *TET1* and *TET2* have been demonstrated by luciferase assays already (28), (29), (30). Interestingly, a different ITC, phenethyl isothiocyanate, was reported to inhibit the upregulation of miR-29b-3p in mouse lung tissue mediated by environmental cigarette smoke (31). The ability of ITCs such as SFN and phenethyl isothiocyanate to modulate miR-29b-3p levels might be at least partially attributable to the induction of Nrf2, which itself has been reported to downregulate miR-29b-3p, via the binding at and consequent repression of transcription from, the miR-29b-1 genetic locus (32). Therefore, experiments to test the ability of miR-29b-3p to bind to these predicted binding sites in the 3’-UTRs of these genes could lead to further insight into SFN’s potential to upregulate tumour suppressor gene expression in colorectal cancer.

In conclusion, SFN upregulates let-7f-5p and let-7g-5p in Caco-2 cells but not in CCD-841. Let-7f-5p is able to bind directly to loci in the 3’-UTRs of CDC25A, HMGA2 and MYC that are computationally predicted as let-7f-5p binding sites, and may thus repress the translation of these oncogenes. The upregulation of let-7f-5p by SFN may thereby contribute to its suppressive effects on colorectal cancer.

